# Mistakes matter, age doesn’t: task outcome modulates implicit motor adaptation similarly in young and older adults

**DOI:** 10.64898/2026.03.11.711139

**Authors:** Matheus Maia Pacheco, Pauline Hermans, Dante Mantini, Alice Nieuwboer, Jean-Jacques Orban de Xivry

## Abstract

Despite several age-related processes impacting motor performance, older adults often retain the ability to implicitly adapt to sensory prediction errors. Here, we leverage the fact that implicit adaptation is not attenuated by aging to study the impact of aging on responses to motor errors. In other domains, such as reinforcement learning, aging has been shown to influence how task outcomes or rewards are processed and used to guide subsequent actions, with some studies emphasizing that older adults react more strongly to a miss than to a hit. We aimed to extend these reinforcement learning findings to the motor domain with two preregistered experiments testing whether missing the target leads to larger implicit adaptation in young and older adults to the same extent. In addition, we compared these results to one reinforcement learning task in the motor domain (Boolean feedback after reaching in the absence of visual feedback) and one in the cognitive domain (reward-based decision making). While we found age-related effects in the cognitive domain, we did not observe a consistent effect of age on the modulation of reaching direction or motor adaptation by task outcomes. These results suggest a domain-specific nature of age-related changes in sensitivity to task outcomes.

## Introduction

During the aging process, individuals experience a range of progressive changes that may reduce their ability to perform optimally in activities of daily life (Ricklefs & Finch, 1995). The literature provides an extensive list of these changes, including decreased strength, slower information processing, reduced attentional resources, and increased muscle force variability (Edström et al., 2007; Enoka et al., 2003; Gopie et al., 2011; Henninger et al., 2010). Considering the ubiquity of such processes in motor activities, the ability to adapt to these changes is therefore of utmost importance.

Sensorimotor adaptation can be defined as a modification in behavior that occurs in response to a sensory discrepancy between expectation and observation (Morehead & Orban de Xivry, 2021). Such discrepancies may result from experimentally induced visual or kinetic changes in the environment or from alterations in body or muscle mechanics (Krakauer et al., 2019). The latter makes the capacity to demonstrate sensorimotor adaptation particularly relevant in aging.

Early studies suggested that older adults adapt less to motor perturbations compared to younger peers (see Heuer & Hegele, 2014; Seidler et al., 2010). However, when the adaptation process is separated into its explicit (intentional) and implicit (automatic) components (see Taylor et al., 2010, 2014; Taylor & Ivry, 2012), it becomes clear that aging affects the explicit component, whereas implicit adaptation appears to be preserved (Hermans et al., 2025; Vandevoorde & Orban de Xivry, 2019, 2020, 2021). Furthermore, when the implicit process is isolated from the explicit one through careful experimental manipulation (Morehead et al., 2017), implicit adaptation seems more pronounced in older adults than in younger ones (Cisneros et al., 2024; de Witte et al., 2026; van de Plas & Orban de Xivry, 2026; Vandevoorde & Orban de Xivry, 2019, 2021).

There are several possible explanations for why implicit adaptation is larger in older individuals (see, for instance, Tsay et al., 2022). A possibility could be that older adults are more sensitive to misses than younger adults and therefore exhibit larger adaptive responses. Numerous studies in younger adults have shown that task outcomes influence the level of implicit adaptation (Al-Fawakhiri et al., 2023; Kim et al., 2019; Leow et al., 2018). This has been demonstrated in the task-irrelevant clamped feedback paradigm (Morehead et al., 2017). In this, the direction of the cursor is rotated by a fixed angle with respect to the target, independently of the actual hand direction, while the radial cursor velocity is matched to that of the hand. Unbeknownst to participants, the discrepancy between cursor motion and the invisible hand trajectory results in a gradual drift of reaching movements in the direction opposite to that of the perturbation (Morehead et al., 2017). With task-irrelevant clamped feedback, one can force the cursor either to hit the target (using a small perturbation angle and a large target) or to miss the target (using the same perturbation angle but a smaller target). Such manipulation of task outcome modulates implicit motor adaptation, as missing the target leads to larger adaptation than hitting it (Kim et al., 2019). Similar results have been obtained using other experimental paradigms (Leow et al., 2018).

Task outcomes such as hitting and missing a target can be considered intrinsic rewards, which strongly modulate sensorimotor processes. For instance, rewards influence a range of mechanisms in motor control, including motor adjustments (Vassiliadis et al., 2021), within-movement compensations (Codol et al., 2023), trial-to-trial correction mechanisms (Palidis et al., 2019), and the shaping of motor variability (Pekny et al., 2015). In the reinforcement learning framework, providing hit or miss information allows one to gradually compensate for a visuomotor rotation (Galea et al., 2015; Izawa & Shadmehr, 2011; Nikooyan & Ahmed, 2015), even though such adaptation appears to rely almost exclusively on explicit processes and re-aiming strategies (Codol et al., 2018; Holland et al., 2018).

Reward-driven changes (and reinforcement learning in general) are influenced by aging (Düzel et al., 2010; Frank et al., 2004). An age-related decline in reward-based reinforcement learning has been observed in the motor domain (Heuer & Hegele, 2014). Beyond motor adaptation, the dopamine hypothesis of cognitive aging posits that older adults exhibit diminished sensitivity to reinforcement, reducing their ability to learn from rewards—particularly negative ones (Van De Vijver et al., 2015). In decision-making tasks where participants must choose between two options, the same dopaminergic decline leads older adults to switch options more often following a negative reward than younger adults (Frank & Kong, 2008; Worthy et al., 2015, 2016; Worthy & Maddox, 2012), making them more sensitive to negative outcomes. In summary, the choice of the next action based on the outcome of the previous one differs between young and older adults. This raises the intriguing possibility that a similar effect may occur in motor adaptation.

Across two pre-registered experiments, we investigate whether task outcome modulates implicit motor adaptation similarly in older and younger adults, and explore the domain- and task-specificity of this effect by testing the same participants in a reward-driven reaching task and a reward-driven decision-making task. The first of these tasks is a controlled reward probability task in which participants receive rewards after a reaching movement depending on an experimentally manipulated probability of reward (Pekny et al., 2015). Following the reinforcement learning literature in motor learning (Sutton & Barto, 2018), a decrease in reward probability leads to an increase in reach direction variability. The second task is a probability learning task in which participants learn to associate different pairs of symbols with different reward probability distributions (Frank et al., 2004; Frank & Kong, 2008; Simon et al., 2010).

We hypothesized that older adults would be more sensitive to misses or negative rewards than younger adults. Therefore, they would exhibit greater implicit adaptation when missing the target (but not when hitting it), show increased reaching angle variability after non-rewarded trials (Pekny et al., 2015), and perform better at avoiding low-reward options rather than selecting highly rewarding ones in the probability learning task (Frank et al., 2004).

## Methods

### Participants

Thirty-two healthy young adults and 30 older adults (60-75 years old) participated in Experiment 1, and 29 healthy young adults and 29 older adults in Experiment 2. They received financial compensation for participating in this study. The inclusion criteria were right handedness (indicated by a score bigger than 40 on the Edinburgh Handedness Inventory), Montreal Cognitive Assessment score above 26, no colorblindness, no mood swings or mood drops, no other disease or disorder that can interfere with the experiment, and no previous experience with motor adaptation tasks. All individuals read and signed an informed consent form to the experimental procedures, which were approved by the ethics committee of KU Leuven (G-2020-2035).

In experiment 2, six participants (one young adult and five old adults) were not included in the analyses provided our exclusion criteria (Montreal Cognitive Assessment score above 26) or not learning the criteria in the probabilistic selection tasks. The resultant number of participants were 29 for each group.

## Equipment and tasks

All data was collected using the KINARM End-Point Lab (KINARM, Ontario, Canada). The equipment is composed of a robotic handle connected to two joints that can move in two dimensions parallel to the ground. Grasping this handle, individuals control a visual cursor presented to them on a mirror positioned above their arm and handle and reflecting the image of a screen on top of the mirror. This setup occluded participants’ view of their hand, ensuring reliance on the screen’s visual feedback. In all tasks, participants controlled the cursor projected on the screen with their right hand.

The two experiments consisted of different versions of the same three tasks: 1) a reaching paradigm with two different tasks (i.e., a task with controlled probability of reward and a task with task-irrelevant clamped feedback), and 2) a probabilistic reward learning paradigm. The order of these two paradigms were counterbalanced across participants. The controlled probability of reward task always preceded the task-irrelevant clamped feedback to avoid any carry-over effect between the adaptation task and the reaching task.

### Experiment 1

#### Task Irrelevant Clamped Feedback

In this task, the task outcome (hit/miss) was controlled, while the sensory prediction error was kept constant (similar to the third experiment of Kim et al., 2019). During baseline, participants made center-out shooting movements from a starting position to small targets (diameter: 3 mm) with veridical feedback (i.e., cursor motion matched hand motion). This was performed for 52 trials on 4 different targets (13 trials per target) located at 45, 135, 225 and 315° from the horizontal axis and 8cm away from the starting position (see Figure 1).

**Figure 1.**
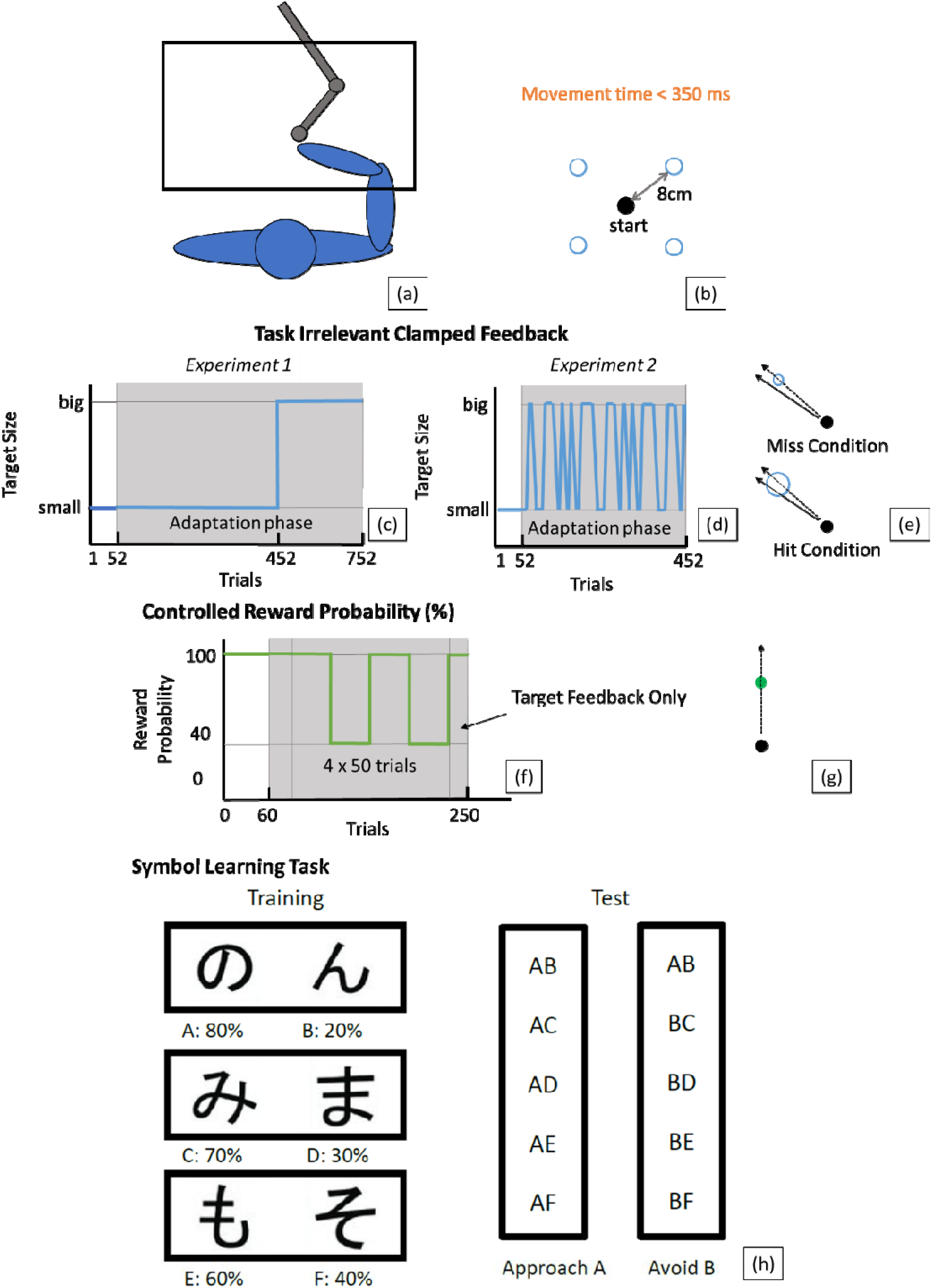
(a) Experimental setting with the individual seated in the KINARM. (b) Visual set-up of the tasks on controlled probability of reward and irrelevant clamped feedback. Participants make reaching movements from a starting point towards a target which are projected on a screen. One blue target per trial was presented (pseudorandom order) in the irrelevant clamped feedback task. Only the green target was presented in the controlled reward probability task. (c) Schedule for the irrelevant clamped feedback task in Experiment 1: individuals started with the vision of their hand in a medium sized target and performed the task for 52trials. After, they switched to the adaptation phase with either to a small or big target for 400 trials where irrelevant concurrent feedback was shown with a deviation of 1.75°. The opposite condition was performed after for 300 trials. (d) Schedule for the irrelevant clamped feedback task in Experiment 2: individuals started with the vision of their hand in a medium sized target and performed the task for 52 trials. After, they switched to the adaptation phase where they reached to four targets (2small and 2 big) for 400 trials where irrelevant concurrent feedback was shown with a deviation of 1.75°. (e) Illustration of the miss and hit conditions: the error feedback was similar; the only difference was the size of the target leading to hits or misses. (f) Schedule for the controlled reward probability task: individuals performed 60 trials to one target with both concurrent feedback of the cursor and target feedback. Then, they performed 25 trials with 100% target feedback only. After this, they performed either the 40% or 100% condition for 50 trials each. (g) Illustration of the condition. (h) Paradigm of the probabilistic selection task. Participants first learned to choose a particular symbol over another with a fixed reward probability for each of the three pairs. In the test session, the participant’s tendency to choose-A over the other symbols and choose any other symbol than B was assessed.

After this baseline phase, the cursor motion was uncoupled from hand motion: while the cursor distance from the home position matched hand distance from the home position, the cursor had a fixed angular offset relative to target position of 1.75°, regardless of the actual hand trajectory. Participants received instructions on the nature of the cursor feedback with a few practice trials (1 target at 0°, randomized clamp direction) and were instructed to ignore this feedback and always try to bring their invisible hand to the visual target. There was no feedback about the actual position of the hand during movement throughout the adaptation phase. Two different targets sizes were used during the experiment. In the Miss condition, a small target was used (diameter: 3 mm), meaning that the 1.75° deviation brought the cursor outside of the target. In the Hit condition a larger target was used (diameter: 8 mm), meaning that the 1.75° deviation brought the cursor inside of the target.

On each trial, a point could be earned with movement durations between 250 and 350ms. Target locations were pseudo-randomly selected in bins of 4 trials. Participants started to reach under clamped feedback for 400 trials with one of the two target sizes followed by 300 with the other target size (700 adaptation trials in total). One-minute breaks were provided after trial 275, 450 and 625. The direction of the error clamp (clockwise or counterclockwise) and the order of the target size (small or big) were counterbalanced across participants within each group.

#### Controlled Probability of Reward

In this reaching task, we aimed to replicate the findings of Pekny et al. (2015) by controlling the probability of reward. Participants made center-out shooting movements from a starting point (diameter: 5 mm) through a medium-sized target (diameter: 5 mm) positioned at a distance of 8 cm at 0° (see Figure 1). In the familiarization phase (60 trials), participants received online feedback. To encourage fast movements, participants received a point whenever a movement lasted between 250 and 350ms. After this baseline, online cursor feedback was removed, being replaced with feedback on trial outcome. Individuals were instructed that the target would turn green for hits and red for misses. However, in this condition, the probability of reward was independent of the actual movement (see Galea et al., 2013) – different from Pekny et al. (2015). The probability of reward was manipulated such that, in the high-reward condition, the participant would receive a reward for 100% of the trials (100% probability of reward) and, in the low-reward condition, the participant received a positive outcome (hit signal) only in 40% of the trials (40% probability of reward). Participants performed 25 trials with 100% reward, then they performed either the 40% or 100% condition for 50 trials each, two times, and then, a last 25 trials were performed with 100% reward (see Figure 1). For each successful movement, the participant received two points. The order of the probability conditions was counterbalanced across participants within each age group.

#### Probabilistic Selection Task

Sensitivity to positive and negative feedback was tested by employing the experimental task of Frank et al. (2004), whereby participants had to learn to the association between pairs of symbols and their probability of reward (Figure 1.g). At the start of the task, the robot guided the participant’s hand to a ‘home’ position, where the participant stayed for the rest of the task. Two symbols were presented on the screen for 5 s. The participant needed to exert an isometric force against the robot handle in the direction of the chosen symbol. In the training phase, three different stimulus pairs (A vs. B, C vs. D, E vs. F) were presented at a random location (left and right) on the screen. Probabilistic feedback was provided to indicate whether the answer given by the participant was rewarded or not. For the easy pair (A vs. B), choosing A had an 80% chance of being rewarded while for stimulus B this was 20%. For the pairs “C vs. D” and “E vs. F”, probabilities were 70/30% and 60/40%, respectively. Difficulty varied between pairs, with pair “E vs. F” being the most difficult to learn. Participants were explained that there was no absolute correct answer and that they would have to identify the stimulus with the highest chance of being correct. Training was completed when participants passed a certain performance criterion for each pair (65% correct responses in “A vs. B”, 60% in “C vs. D”, and 50% in “E vs. F”), which was evaluated every 60 trials, with a maximum of six training rounds. In the test phase (90 trials), choice behavior was tested by creating comparisons between all possible figures in a randomized order (e.g., “A vs. D”, “B vs. D”, “A vs. B”). Each test pair was presented 6 times.

After the whole data collection, we found that the probabilities of the pairs were assigned incorrectly. That is, all low-probability symbols (B, D and F) were assigned a probability of reward of 0%. Thus, even though the pairs still might preserve the hierarchy (e.g., “A” being more probable than other symbols), we cannot ensure that we can replicate the results of this task (Frank et al., 2004; Frank & Kong, 2008; Simon et al., 2010). The correct version of the task was used in experiment 2.

### Experiment 2

Three modifications were implemented. First, we modified the task irrelevant clamped feedback as to preclude the need for controlling for the order of targets (see below). Second, in the Controlled Probability of Reward task, we emphasized the fact that the target did not change its position. This was done to minimize potential explicit exploration of different target positions (angles) whenever the reward probability decreased. Third, we corrected the technical issues on assigning probabilities to the low-probability symbols in the probabilistic learning task and matched the accuracy estimates to those of Frank et al. (2004).

#### Task Irrelevant Clamped Feedback

In this version of the task, the baseline was the same as in Experiment 1: center-out reaching movements with veridical cursor feedback (52 trials, 4 different target locations at 45, 135, 225 and 315°). After this baseline, the adaptation phase was similar to Experiment 1. The difference is that a pair of the targets was always large while the other pair was always small. Each pair would be either at the right (45° and 315°) or left (135° and 225°) from the center position of the hands (or above/ below, counterbalanced between participants). The direction of the clamped irrelevant feedback was always ± 1.75° from the direction of the target (being positive or negative counterbalanced between participants). The participants performed a single adaptation condition for 400 trials.

### Data Processing and Statistical Analyses

#### Pre-Registration

Each experiment was preregistered online at Open Science Foundation (Experiment 1: osf.io/3syz5/; Experiment 2: osf.io/hjm76) containing the main hypotheses, experiments, key dependent measures and statistical comparisons. The preregistration considered 30 participants per group, but two extra healthy young adults were collected in Experiment 1 given their availability. Two participants (one healthy young adult and one older adult) in Experiment 2 failed to pass the minimum performance in the symbol learning task and were, therefore, excluded from analyses. The preregistered comparisons are the numbered analyses described below while exploratory analyses are presented in the results sections. Slight modifications in the statistical tests were performed from pre-registered Experiment 1 as the majority of experiments demonstrated deviations from normality. Instead of using the preregistered non-parametric analyses (Friedman’s ANOVA) which would limit the inclusion of possible interactions, we preferred to perform robust analyses (Wilcox, 2022). Experiment 2 pre-registration already considered robust analyses.

The number of trials of each task deviated slightly from the pre-registered report of Experiment 1. For the controlled probability task, the pre-registration reported 50 trials of baseline while the actual data collection had 60 trials. Additionally, the pre-registration stated that the four blocks of either 100% or 40% probability of reward would start immediately after baseline. In the present experiment, the blocks started after 25 trials in 100% reward condition to familiarize participants with the condition and ended with another 25 trials in 100% reward condition. For the clamped irrelevant feedback task, the pre-registration file stated that individuals would perform 50 trials; to have the same number of trials for each target, we changed the number of trials to 52. The pre-registration file also stated that 1-min breaks would be given after trials 525, 700 and 875. These trial numbers reflect the clamped irrelevant feedback task trials added with 250 trials supposed to occur in the controlled reward task. Finally, the Experiment 1 pre-registration also stated that individuals would perform an extra experiment of smartphone-based tapping task that would be collected before, between the reaching tasks and probability learning task, and at the end of the session as a control task – without any strong hypothesis. However, provided technical issues (non-availability of the equipment when collecting data of some participants), not all participants were tested in this paradigm. Thus, the data is not discussed or analyzed here.

For both experiments, the pre-registration files explicitly stated the hypotheses and, thus, the expected effects from the analyses. However, as we were interested in evaluating whether other results (specifically main effects) were to occur, we employed a correction to limit Type I error in our analyses – see below. This was not planned in the pre-registration of Experiment 1

#### Data

For the task irrelevant clamped feedback and controlled probability of reward, the primary dependent variable is the hand angle relative to the target at 4 cm into the movement (Vandevoorde & Orban de Xivry, 2019). Hand angles were calculated and corrected for baseline error by subtracting the average error of the last 20 baseline trials from all parameters. Trials with angular errors greater than 60° were considered outliers and removed from further analysis (≈ 0.16% of trials were removed in Experiment 1). For probabilistic selection task, the main outcomes were overall accuracy in given conditions that were calculated on the basis of proportion of correct responses given all trials in that given condition.

Data were extracted from KINARM files in MATLAB 2020b (MathWorks) scripts designed for this purpose. The statistical analyses (described below) were performed in either RStudio 1.3.1093 or JASP 0.14.1. The functions for robust analyses were extracted through Rallfun-v38.txt (the most updated list of functions from Wilcox, 2022). All scripts are openly available at https://osf.io/rjfwq/. Effect sizes (i.e., Cohen’s *d* for *t*-tests, η ^2^ for regular ANOVA and ξ for robust analyses) were reported and when interaction effects occurred, the post hoc analyses refer to the interaction effects. *d* values above 0.2, 0.5 and 0.8, η ^2^ values above 0.01, 0.06, and 0.14 and ξ values above 0.15, 0.35, and 0.50 are considered small, medium and large effects, respectively (Richardson, 2011; Wilcox, 2022). Results are reported either in terms of means ± standard deviation (i.e., M ± SD) or median and interquartile range (i.e., Med, IQR).

For three-way robust ANOVAs considering a three-way interaction with two between-subject and one within-subject effect, there is, as far as we are aware, no described way to perform post hoc analyses and to compute ξ effect size. In these cases, we performed robust simple main effects with *yuenv2* and *yuendv2* (for independent and dependent effects, respectively) using Holm’s sequential post hoc procedure (Abdi, 2010). In these cases, the effect sizes are provided in terms of the simple main effects. When a two-way interaction was the highest-order interaction found in three-way robust ANOVAs, we pooled the data and considered it as a two-way robust ANOVA to calculate effect sizes and perform the post hoc analyses.

For all analyses, we corrected for the multiplicity of ANOVA by controlling for the false discovery rate as described in Cramer et al. (2016). For this, we used the *p.adjust* function in *R* using the Benjamini Hochberg correction. We decided to make this correction despite our pre-registration of the expected effects (see above).

#### Experiment 1: Analysis 1.1: Implicit adaptation level with task irrelevant clamped feedback

For this analysis, the main outcome was the average hand angle during trials 300 to 400 of the first adaptation block (see Figure 1) corrected for baseline bias. In the pre-registration file, we stated that based on robust results in previous studies measuring implicit adaptation with task irrelevant clamped feedback (Morehead et al., 2017; Vandevoorde & Orban de Xivry, 2019), participants who move in the opposite direction of the expected one (average hand angle of the last 100 trials < 0°), have ignored the instruction (i.e. moved their hand in the direction of the error clamped cursor instead of towards the target) and have their data thus discarded. However, no participant showed this behavior.

Hand angle was compared in a robust two-way ANOVA with age group (young, older) and task condition (hit, miss) as between-subjects factors. We expect to find a main effect of task outcome such that participants reach higher adaptation levels while missing the small target than while hitting the bigger target. Furthermore, we expect to find a main effect of age group with older adults showing higher implicit adaptation levels than their younger controls. Lastly, we expect to find an interaction effect indicating that older adults only show higher adaptation levels than young adults in the miss condition as this would demonstrate a higher sensitivity in terms of no-reward conditions.

#### Analysis 1.2: Adaptation level modulated by task outcome with irrelevant clamped feedback

The average hand angle for both target sizes was calculated and corrected for baseline error as described in Analysis 1 (trials 300 to 400 of the first adaptation block and trials 200 – 300 of the second adaptation block, see Figure 1). These hand angles were compared in a three-way repeated measures robust ANOVA with age group (young, older) and order of conditions (first hit, first miss) as between subject factors and task condition (hit, miss) as within-subjects factor. We expect to find an interaction between conditions and order of conditions where the first condition will determine the increase in adaptation of the second and an interaction between age and condition given the highest tendency of older adults to be more sensitive to no-reward conditions.

#### Analysis 1.3: Within-subject variability in hand angle with controlled reward probability

In controlled probability of reward task, the primary dependent variable is the variability in trial-to-trial changes in reaching hand angle – calculated as follows. Change in hand angle (u) from trial n to trial n+ 1 followed the following formula:

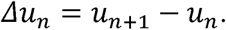

Thus, we calculated the standard deviation of Δ*u* for rewarded and unrewarded trials separately. Standard deviation (*SD_U_*) was compared with a repeated measures robust ANOVA with reward condition (rewarded, unrewarded) as within-subjects factor and age group (young, older) and order of reward condition (40%, 100% first) as between-subjects factors. We expect an increase in trial-to-trial hand angle variability after unrewarded trials compared to the rewarded trials (main effect of condition). Furthermore, older adults should show a larger variability in the unrewarded trials than younger adults (an interaction between condition and age).

We considered that the *SD_U_* from the 40% reward condition would combine the effect of both rewarded and non-rewarded trials. For this reason, as an exploratory analysis (not pre-registered), we also compared the variability in reaching angle (*SD_U_*) between age groups as a function of reward within the 40% reward condition (i.e., compared rewarded and non-rewarded trials).

#### Analysis 1.4: Correlation between the modulation of adaptation and modulation of variability in hand angle given outcome

Modulation of adaptation level by task outcome was calculated by subtracting the adaptation level in the hit condition from the adaptation level in the miss condition (heretofore MA, see Analysis 1.2 for the calculation of adaptation level). The variability in hand angle exhibited at 100% reward condition was subtracted from the one at the 40% reward condition (heretofore Δ*SDu*, see Analysis 1.3 for the calculation of variability). MA was estimated using a robust linear regression with the following independent variables: Δ*SDu*, a binary age vector (Age), a binary order vector (0) and the interactions of Δ*SDu* and Age (the correlation between MA and Δ*SDu* could be different across age groups) and of Age and 0 (the order effect might be different as a function of age group). Each of these factors were indexed with a coefficient (b, c, d, e, and f) and an intercept a:

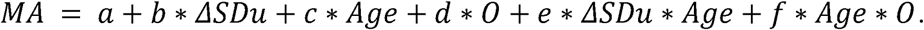

We expect to find a correlation between Δ*SDu* and MA, which would be associated with a significant coefficient b. Both tasks measure people’s response to an unsuccessful movement as compared to a successful one. Participants who show a higher modulation in adaptation rate to task outcome will have a greater increase in hand angle variability after unrewarded trials.

#### Analysis 1.5: Probabilistic selection

The primary dependent variable is accuracy in the test phase in terms of choosing A whenever it appeared in a pair and avoiding B whenever it appeared in a pair. We looked at the accuracy in choosing symbol A (proportion of trials in which A was chosen over the other five possible symbols) and avoiding B (proportion of trials in which the other five possible symbols were chosen over B). Accuracy of approaching A corresponds to learning based on positive feedback, while accuracy of avoiding B corresponds to learning based on negative feedback. These accuracies were compared in a robust repeated measures ANOVA with subject group (young – older) as between-subjects factor and type of condition (choose-A, avoid-B) as within-subjects factor. We expect to find an interaction between the difference in approach/avoid behavior and age group (i.e., an interaction between conditions and age). The difference in accuracy between the age groups would be larger for the negative feedback condition than the positive feedback condition, with older adults learning more from negative feedback.

#### Analysis 1.6: Correlation between the modulation of the adaptation and selection modulation given outcomes

We subtracted the accuracy on avoid-B from choose-A to measure the difference on deciding on rewarding choices and avoiding no-reward (AB, see Analysis 1.5 for calculation of accuracy). MA (see Analysis 1.4) was estimated using a robust linear regression with AB, a binary age vector (Age), a binary order vector (0) and the interactions of AB and Age (the correlation between MA and AB could be different across age groups) and of Age and 0 (the order effect might be different in function of age group). Each of these factors were indexed with a coefficient (b, c, d, e, and f) and an intercept a:

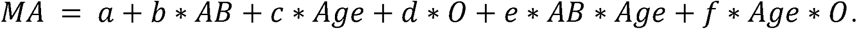

We expect to find a correlation between MA and AB, which would be evidenced by a significant coefficient b. More specifically, participants who show more adaptation after a miss relative to a hit will learn more easily from negative feedback relative to positive feedback.

**Experiment 2:** In the second experiment, provided the modifications (see above), we modified the clamped irrelevant feedback task from Experiment 1, avoiding the order effect observed in the results. Additionally, provided exploratory analyses in Experiment 1, we found that the controlled probability of reward task would be better analyzed from variability of change (SD*u*) within the 40% reward condition in terms of rewarded and non-rewarded trials instead of comparing the 100% and 40% reward conditions.

#### Analysis 2.1: Implicit adaptation level with task irrelevant clamped feedback

This Analysis followed the same rationale as the Analysis 1.1 in Experiment 1. That is, we corrected the adaptation outcomes (average hand angle in the last 100 trials) in terms of the baseline controlling it in terms of which pairs corresponded to hits or misses. Then, the hand angle was compared in a robust two-way ANOVA with age group (young, older) as between-subjects factor and task condition (hit, miss) as within-subjects factors. We expect to find a main effect of task outcome such that participants reach higher adaptation levels while missing the small target than while hitting the larger target. Mainly, we expect to find an interaction effect indicating that older adults show different adaptation levels than young adults in the miss condition as this would demonstrate a differential sensitivity in terms of no-reward conditions.

***Analyses 2.2 to 2.5*** followed the same rationale as Analyses 1.3 to 1.6, respectively. Analyses 2.2 and 2.3 modified the dependent variable (planned in the pre-registration of Experiment 2): SD*u* within the 40% reward condition in terms of rewarded and non-rewarded trials instead of comparing the 100% and 40% reward conditions. Analyses 2.4 and 2.5 were identical to analyses 1.5 and 1.6.

As an extra analysis (not pre-registered), we also calculated the Cohen’s *d* effect size and its 95% confidence interval for the results of the main results of the three paradigms. For the task irrelevant clamped feedback, the main result is the interaction between age and hit vs. miss. For this, we calculated the interaction effects for (a) those who started with the miss condition in the first experiment, (b) those who started with the hit condition in the first experiment, and (c) all participants in the second experiment. Then, we performed a meta-analysis to summarize these into a single effect size (following Carter & McCullough, 2014 procedure). For the controlled reward probability task, we calculated the interaction effects of age and rewarded trials (within the 40% reward condition) considering all participants of both experiments. For the probabilistic learning task, we calculated the interaction effects of age and approach/avoidance considering the participants of the second experiment. Provided the data fails to follow parametric assumptions, the need to perform transformations between ξ and *d* to have the interaction in terms of *d*, and the lack of a simple transformation between these two measures, we performed a bootstrap procedure (2000 iterations) to calculate each η *^2^* and then transformed its mean to *d* following Cohen (1988) procedure.

## Results

### Experiment 1

#### Age does not affect implicit motor adaptation in the task irrelevant clamped feedback

In the task irrelevant clamped feedback, we investigated whether older individuals would be more influenced by missing the target than younger peers. Regardless of their group and condition, all participants exhibited a gradual drift in the direction of their reaching movements when the visual feedback was perturbed (Figures 2a and 2b). For both age groups, the task outcome appeared to modulate the level of adaptation in both the first and second phase of the adaptation period. At the end of baseline, participants from both groups showed similar hand angles around the target (mean ± standard deviation; young: 0.13 ± 1.06°; older: −0.14 ± 2.34°; *t* [60] = 0.58, *p* = .566, *d* = 0.15). At the end of the first adaptation block, missing the target led to larger adaptation levels (Analysis 1.1; young: 17.95 ± 5.96, older: 16.29 ± 7.42) than hitting the target (Figures 2c and 2d; young: 11.77 ± 5.91, older: 11.28 ± 6.28; *Q* = 15.24; *p* = .003; ξ = 0.68). Contrary to our expectations, we found no evidence that the influence of task outcome on the adaptation response differed between young and older adults (interaction between age and condition: *Q* = 0.13; *p* = .722; ξ = 0.11) or in any condition (main effect of age: *Q* = 0.40; *p* =.722; ξ = 0.14).

**Figure 2.**
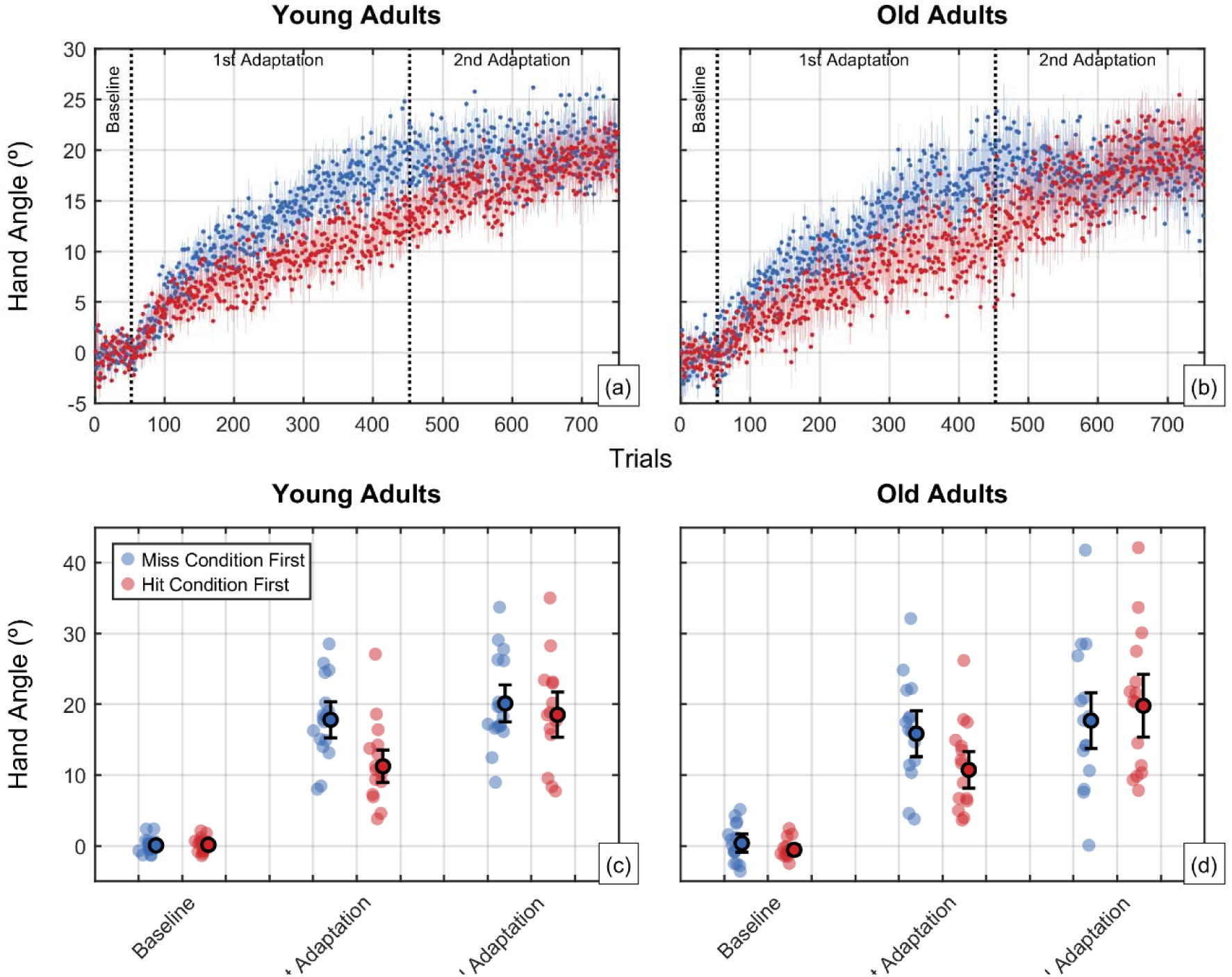
(a) and (b) Hand angle per trial as a function of order of presentation (red: hit first; blue: miss first) for young and old adults, respectively, in the irrelevant clamped feedback task. The dots and shaded area represent the mean and standard deviation. (c) and (d) Winsorized mean (dark border circles), 95% confidence interval (error bars) and distribution (colored circles) of the hand angles per adaptation stage of the task.

These effects are maintained when considering the second block of the adaptation period where target sizes were switched. In that period, older and younger adults show similar adaptation levels (Analysis 1.2; main effect of age: *Q* = 0.30; *p* = .870) and participants adapted more in the miss than in the hit condition (main effect of condition: *Q* = 29.01; *p* < 0.001). As expected, the order in which the hit and miss conditions were presented influenced the adaptation levels (interaction condition and order: *Q* = 58.38; *p* < .001; ξ = 0.62). Individuals showed larger adaptation levels initially if they started in the hit condition than if they started in the miss condition (*p* < 0.001), but this difference disappeared after the change in target size. Those who started in the miss condition showed adaptation levels of 17.96 ± 5.69° (young) and 16.29 ± 7.42° (older) with a significant increase to 20.20 ± 6.38° (young) and 18.09 ± 10.37° (older) when in the hit condition (*p* = .006). Starting with the hit condition led to adaptation levels of 11.77 ± 5.91° (young) and 11.29 ± 6.29° (older) with a significant increase to 18.95 ± 7.30° (young) and 20.27 ± 10.04° (older) when in the miss condition (*p* < .001). In summary, older and younger adults appeared to module their adaptation levels as a function of task outcome (missing or hitting the target) similarly in an irrelevant clamped feedback condition.

#### Older adults show less reward-dependent variability than young adults

In the controlled reward probability task, we questioned whether older adults would demonstrate large variability in reaching angle when the probability of reward was decreased compared to young peers. We found that, as expected, participants showed larger variability in reaching angle in the 40% reward condition compared to the 100% reward condition (main effect of condition: *Q* = 17.35; *p* < .001; ξ = 0.30). Surprisingly, older adults showed less variability in reaching angle than their young controls (main effect of age: *Q* = 11.11; *p* = .005; ξ = 0.47) but, contrary to our expectations, this was not specific to the 40% reward condition (Figure 3a; interaction between age and condition: *Q* = 0.67; *p* = .492; ξ = 0.07). Older adults showed (mean ± standard deviation) a variability of 3.76 ± 1.46° and 3.14 ± 0.82° in the 40% and 100% reward conditions while young adults showed 4.77 ± 2.88° and 3.79 ± 1.17°.

**Figure 3.**
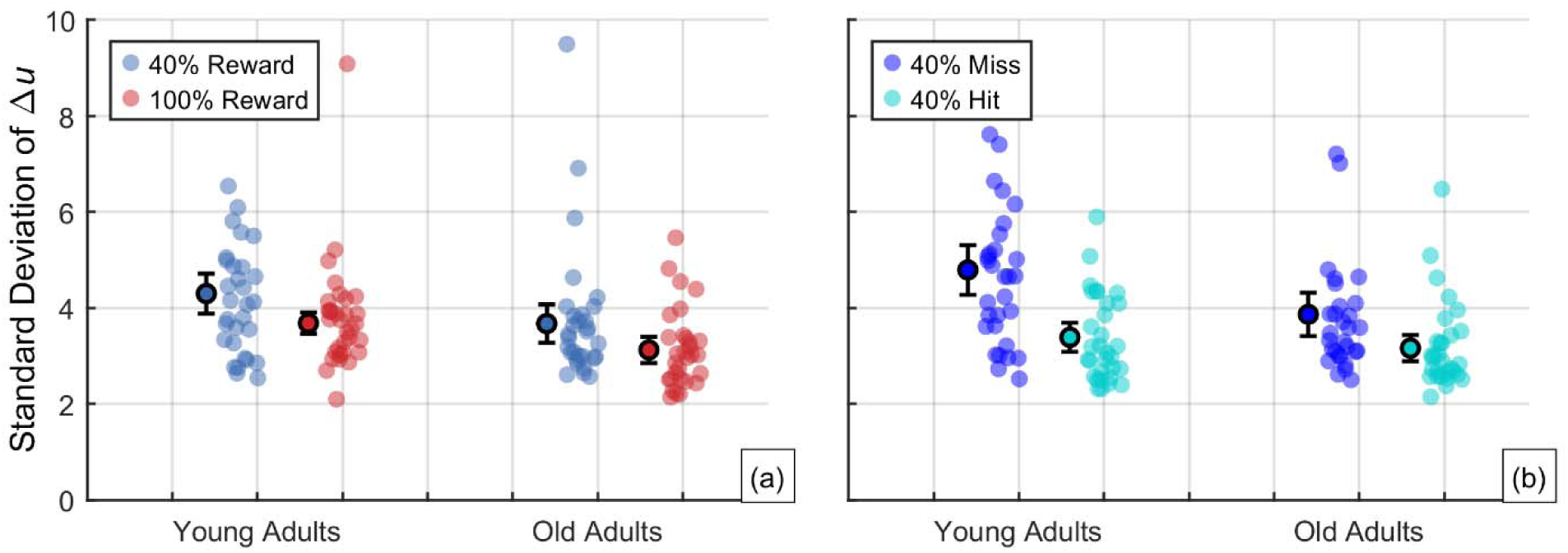
Winsorized mean (dark border circles), 95% confidence interval (error bars) and distribution (colored circles) of the (a) Standard deviation of the trial-to-trial change (Δ*u*) as a function of condition and age group and (b) standard deviation of the trial-to-trial change (Δ*u*) considering rewarded and non-rewarded trials within the 40% probability of reward condition.

Provided the 40% reward condition combines the effect of both rewarded and non-rewarded trials, we also compared the variability in reaching angle between age groups as a function of reward within the 40% reward condition (not pre-registered for Experiment 1). Both young and older adults showed a decrease in reaching angle variability from non-rewarded (young: 5.35 ± 3.45°; older: 4.00 ± 1.79°) to rewarded trials (young: 3.65 ± 1.89; older: 3.20 ± 0.92; *Q* = 50.27; *p* < .001; ξ = 0.54). Older adults were less variable in reaching angle after no-reward trials than young adults (interaction between trial type and age: *Q* = 7.47; *p* = .015; ξ = 0.34; post hoc: *p* = .007) but not in the reward trials (*p* = .196) (Figure 3b).

In summary, older adults showed less variability in reaching angle than young adults. This was mainly related to a low increase in variability when rewards were less frequent (shown by a significant difference in hit/miss comparison within the 40% reward condition).

#### Task-irrelevant clamped feedback adaptations fail to relate to reward-induced variability

To understand whether task implicit adaptations to clamped feedback conditions are driven by the same reward-induced processes as reward-induced increases in variability, we tested whether the difference in implicit adaptation was modulated by the hit/miss conditions (in the task implicit clamped feedback) with the increase in reaching angle variability modulated by the decreasing reward probability (difference in reaching angle variability between 100% and 40% conditions in the controlled reward probability Task) (Analysis 1.4). We did not find evidence that implicit adaptation and reward-induced variability were related to each other (*p* = .659). As in Analysis 1.1, only the order of presentation (in the task-irrelevant clamped feedback) was related to adaptation level modulated by hit/miss conditions (normalized estimate = 1.45; CI_95%_ = [1.24, 1.91]; *p* < .001). These results were maintained if we considered the increase in reaching angle variability modulated by non-rewarded trials within the 40% reward probability condition.

### Experiment 2

#### Age does not affect the influence of task outcome on implicit motor adaptation

In this new version of the task-irrelevant clamped feedback task, participants adapted to the same perturbation as in Experiment 1 but the target sizes varied across targets. During the adaptation period, participants reached to four different targets: two were small (miss condition) and two were larger (hit condition). Figures 4a and 4b show the hand angle of young and older adults as a function of target type during this task at the end of the adaptation phase (Analysis 2.1). Considering the last 20 trials of the adaptation phase, participants of both groups showed similar hand angles around both targets (main effect of age: *Q* = 2.17: *p* = .224; ξ = 0.20) while showing smaller adaptation for hit (young: 13.03 ± 5.40°; old: 10.47 ± 6.15°) than miss conditions (young: 20.68 ± 8.52°; old: 18.88 ± 7.21°) (*Q* = 57.74; *p* < .001; ξ = 0.84). Contrary to our expectations, we did not find evidence that the influence of task outcome (miss vs. hit) on adaptation levels differed between young and older adults (interaction age and target: *Q* = 0.01; *p* = .908; ξ = 0.14).

**Figure 4.**
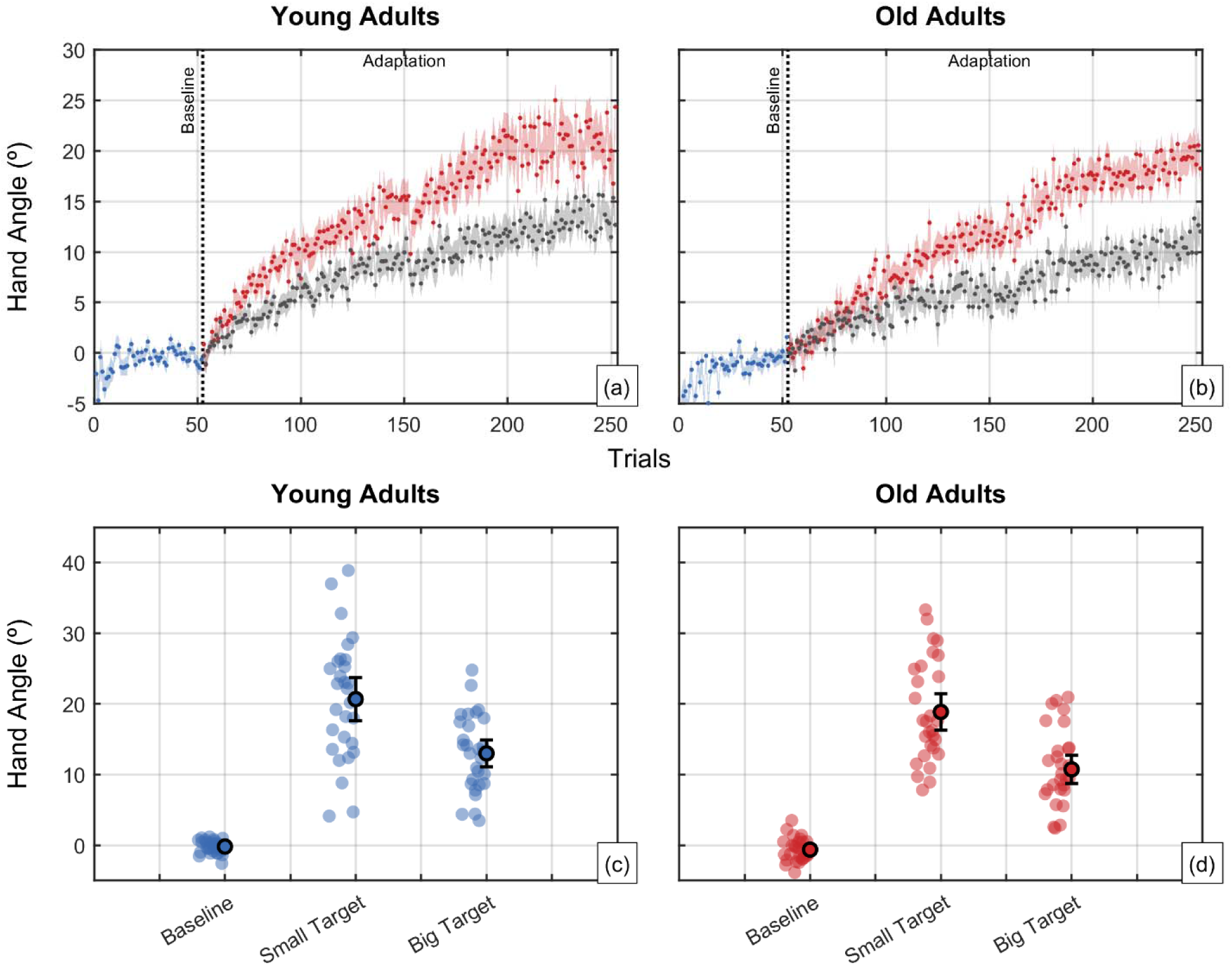
(a) and (b) Hand angle per trial as a function of order of presentation (blue: baseline; red: small targets [miss]; gray: large targets [hit]) for young and old adults, respectively, in the irrelevant clamped feedback task. The dots and shaded area represent the mean and standard deviation. (c) and (d) Winsorized mean (dark border circles), 95% confidence interval (error bars) and distribution (colored circles) of hand angles per adaptation stage of the task.

#### Age does not affect reward-dependent variability

In the controlled reward probability task, we questioned whether older adults would demonstrate large reaching angle variability when the probability of reward was decreased compared to younger peers (Figure 5). Comparing the reaching angle variability between age groups as a function of reward within the 40% reward condition (Analysis 2.2), we did not find, contrary to the previous experiment and expectations, that young and older adults show any difference in trial-to-trial variability in rewarded (young: 4.37 ± 2.05°; old: 4.01 ± 1.50°) and non-rewarded trials (young: 5.07 ± 3.89°; old: 4.69 ± 1.78°) (main effect of age: *Q* < 0.01; *p* = .976; ξ < 0.01; main effect of reward: *Q* = 4.87; *p* = .077; ξ = 0.47; interaction age and reward: *Q* = 4.14; *p* = .077; ξ = 0.24).

**Figure 5.**
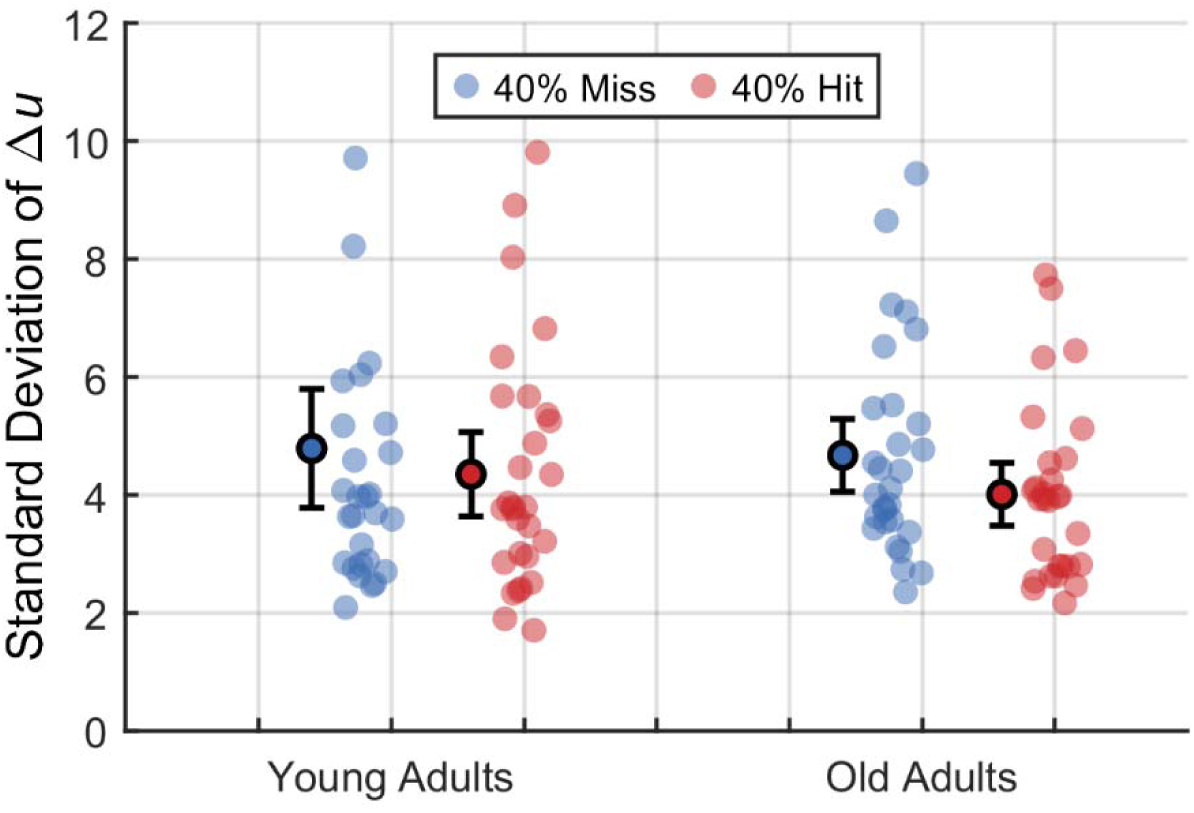
Winsorized mean (dark border circles), 95% confidence interval (error bars) and distribution (colored circles) for the standard deviation of the trial-to-trial change (Δ*u*) considering rewarded and non-rewarded trials within the 40% probability of reward condition.

#### Task implicit clamped feedback adaptations relate to reward-induced variability

To understand whether task implicit adaptations to clamped feedback conditions are driven by the same reward-induced processes as reward-induced increases in variability, we performed a robust correlation between the difference in adaptation level between hit and miss conditions (in the task implicit clamped feedback) with the increase in reaching angle variability from rewarded and non-rewarded trials in the 40% reward probability condition (in the controlled-probability task) (Analysis 2.3).

In line with our previous experiment, we did not find any evidence that those who increased more their reaching angle variability given non-rewarded trials did not show more adaptation in the task implicit clamped feedback in the miss condition compared to the hit condition (normalized estimate = −0.06; CI_95%_ = [−0.35, 0.93]; *p* = 0.453). Nonetheless, we found an interaction effect between age and reaching angle variability (normalized estimate = −0.70; CI_95%_ = [−1.15, −0.05]; *p* = 0.029). To untangle the interaction, we performed two regression analyses, one per age group. The results showed that only young adults increased their adaptation in the task implicit clamped feedback as a function of reaching angle variability (normalized estimate = 1.22; CI_95%_ = [0.36, 1.91]; *p* < .006); older adults did not show such a relation (normalized estimate = −0.76; CI_95%_ = [−1.45, 0.41]; *p* = .388).

#### Older adults avoid less non-rewarding choices than young adults in probabilistic learning

Figure 6 shows the accuracy of young and older adults in learning to choose rewarding options and learning to avoid non-rewarding options in a symbol learning task (Analysis 2.4). While older adults were less accurate in avoiding non-rewarding options than choosing rewarding ones (interaction effect between age and conditions: *Q =* 6.00; *p* = .030; ξ = 0.40; choosing_older_ = 0.79 ± 0.29; avoiding_older_ = 0.58 ± 0.32, *p* = .013), young adults were equally accurate between conditions (choosing_young_ = 0.80 ± 0.20; avoiding_young_ = 0.81 ± 0.30, *p* = .999). Young and older adults differed only when avoiding non-rewarding options (*p* = .015) and not when choosing the rewarding one (*p* = .383).

**Figure 6.**
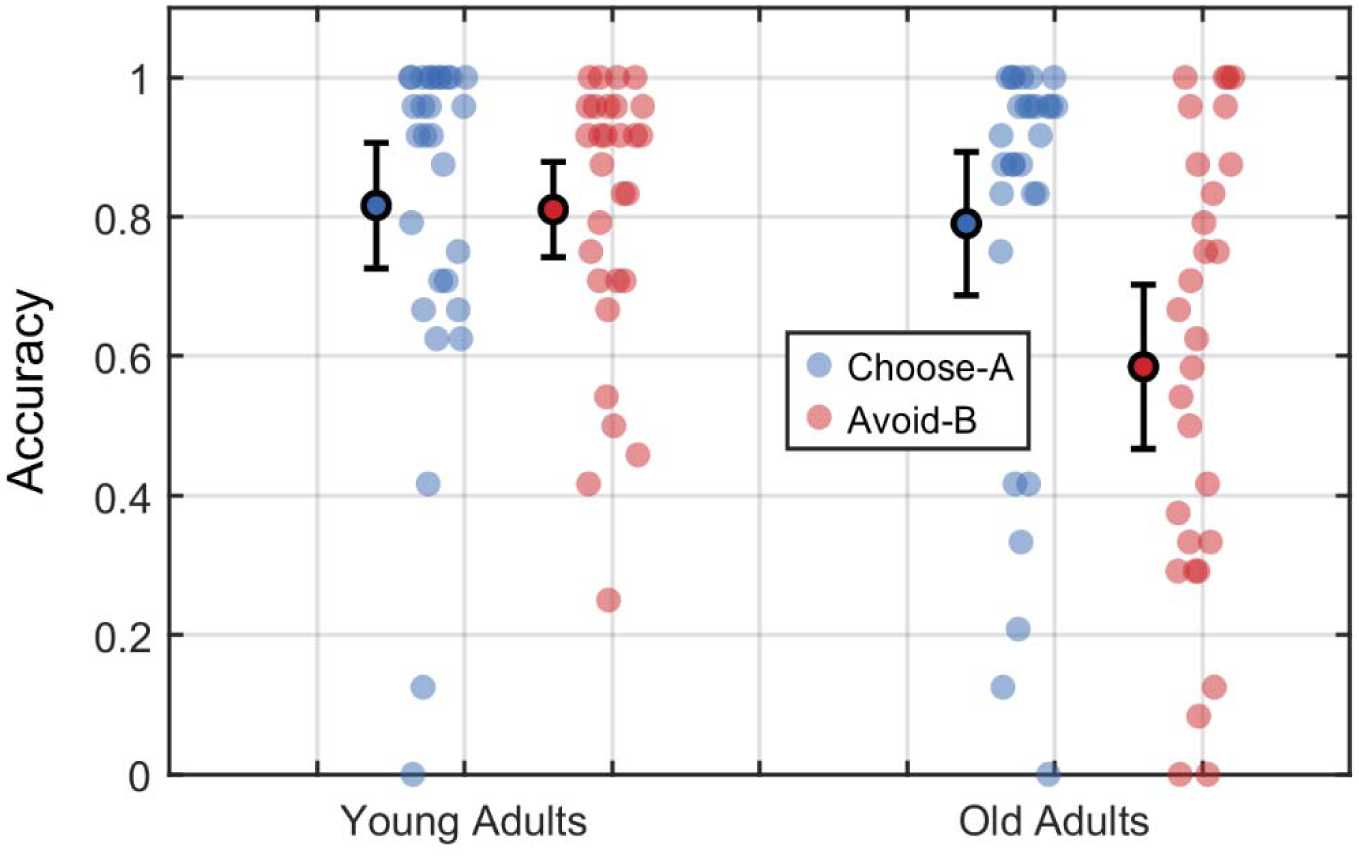
Winsorized mean (dark border circles), 95% confidence interval (error bars) and distribution (colored circles) for the accuracy of young and old adults in choosing-A and avoiding-B in the probabilistic learning task.

#### Task implicit clamped feedback adaptation fails to relate to probability learning approach/avoidance behavior

To understand whether task implicit adaptations to clamped feedback conditions were driven by the same reward-induced processes as in choosing rewarding options and avoiding non-rewarding ones, we performed a robust regression between the adaptation modulated by the hit/miss conditions (in the task implicit clamped feedback) with the difference in the accuracy between avoiding non-rewarding options and choose rewarding ones (in the probabilistic learning task) (Analysis 2.5). The correlation was not observed (*AB*: normalized estimate = −1.38; CI_95%_ = [−5.73; 5.49]; *p* = .741; age: normalized estimate = −0.69; CI_95%_ = [−2.61; 1.80]; *p* = .695; interaction: normalized estimate = 0.48; CI_95%_ = [−6.01; 5.38]; *p* = .904).

#### Summary Results

To summarize the results of the present study, we performed a bootstrapped Cohen’s *d* calculation (2000 iterations) comparing the interacting effects of age and the main outcomes of each task (e.g., miss trials adaptation for task irrelevant clamped feedback) (Figure 7). As it is observed, for the first and second paradigms, the confidence interval encompasses the null hypothesis (*d* = 0). The only clear difference was the interaction effect for probability learning.

**Figure 7.**
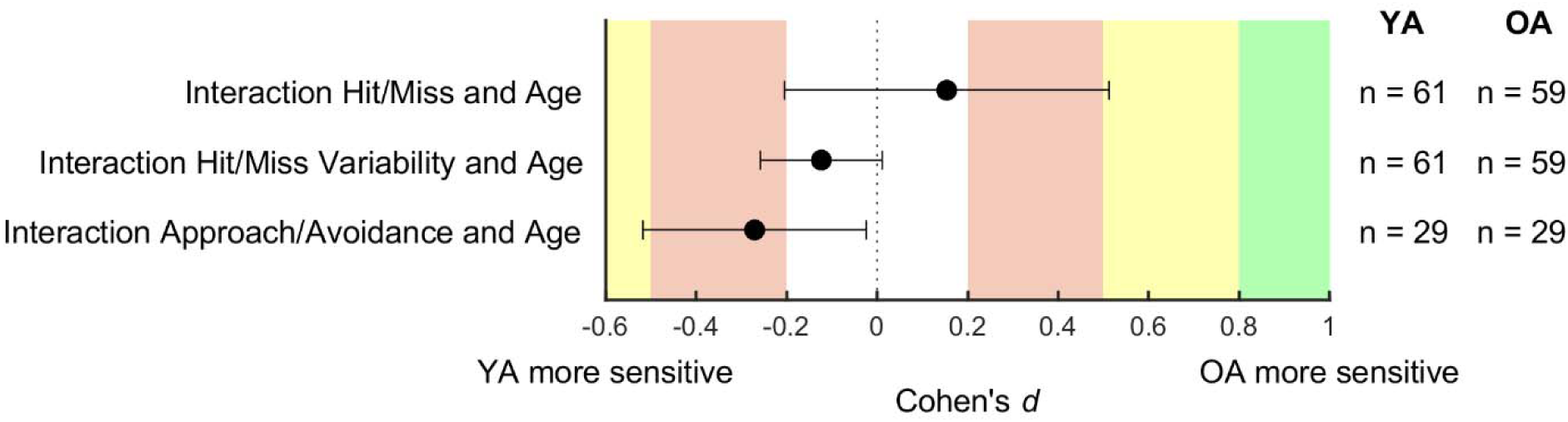
Cohen’s *d* effect size for comparisons between young and older adults pooled for both experiments (reward learning and motor adaptation) and for experiment 2 (probabilistic learning).

## Discussion

In this study, we investigated whether the modulation of behavior by task outcomes is similarly affected by aging across tasks in the motor and cognitive domains. We found that task outcomes modulated motor adaptation similarly for young and older adults. The modulation of reach variability by the presence or absence of rewards was clear in Experiment 1 but more limited in Experiment 2, where re-aiming strategies were minimized. The dependence of reach variability on reward was smaller in older adults than in younger ones, but this effect disappeared when re-aiming strategies were minimized in Experiment 2. Finally, older adults were less accurate in avoiding low-reward choices (Experiment 2). Together, these results suggest that aging affects outcome-dependent processes differently as a function of domain.

### Task outcome modulates implicit motor adaptation similarly in young and older adults

The main goal of this study was to test whether implicit sensory motor adaptation in aging would be differentially affected by task outcome (in terms of reward-driven processes). Our results were unanimous in pointing out that implicit sensory motor adaptation is influenced by task outcome, but that this effect is independent of age. Kim et al. (2019) and Al-Fawakhiri et al. (2023) demonstrated that, using the same task-irrelevant clamped feedback paradigm, implicit sensorimotor adaptation is attenuated when the cursor hits the target compared to when it misses it.

The effect of aging on reward-dependent processes in the motor domain is a complex topic that has garnered considerable attention in recent research. Similarly to our results, the modulation of movement vigor by reward-dependent processes appears to be unaffected by age (Tecilla et al., 2023). Reaction time is also similarly modulated by reward feedback in young and older adults, although the reaction time itself is largely impacted by age (Drueke et al., 2015). In a sequential reaching task, reward similarly modulated learning in young and older adults (Aves et al., 2021). Other studies found more mixed results. For instance, positive reinforcement similarly modulates total adaptation in young and older adults, while negative reinforcement had no effect on the rate of adaptation in both groups (Huang et al., 2018). In contrast, retention was reduced by negative reinforcement in young adults only (Huang et al., 2018). However, these studies are only relevant insofar as missing and hitting the target are considered as negative and positive reinforcement or feedback. Yet, as highlighted by the studies above in the motor domain and consistent with our results, there seems to be little age-related effect on the modulation of motor behavior by either rewards or task outcomes.

### Reward-driven differences between young and older adults are task-dependent

One of the assumptions of the present study was that young and older adults are differentially affected by reward (reinforcement). We also considered that such an effect could be task-specific and, for this reason, considered two classic examples of tasks that have been used in literature to study reward or outcome dependent processes. Considering reward-dependent variability in a motor task, we found inconsistent results between the two experiments.

Considering the exploration-exploitation trade-off in motor control, it is believed that failing to get the expected reward leads to an increase in movement variability to explore the motor space (Dhawale et al., 2017; Wu et al., 2014) and find new motor solutions. In the framework of the Pekny study and in the absence of any visual information about the actual performance, failing to get a reward leads to an increase in reach variability (Pekny et al., 2015). While we could replicate these effects, they were more pronounced in experiment 1 than in experiment 2. It is possible that some of these effects are driven by conscious re-aiming strategies as those that are developed in reinforcement learning experiments (Codol et al., 2018; Holland et al., 2018). Future studies are necessary to investigate the effect of aging on reward-dependent variability of reaching movements and whether these effects are linked to re-aiming strategies.

### Are age effects absent in the motor domain but present in the cognitive domain?

While the modulation of behavior by reward or outcome in the motor domain was not affected by age, there were age-related effects in the probabilistic learning task. Given that increased reach variability due to the absence of reward and learning reward contingency are both affected by Parkinson’s Disease (Frank et al., 2004; Karrer et al., 2017; Pekny et al., 2015), we anticipated that performance in these two tasks would be similarly affected by aging but this is not what we found. While aging did not seem to impair the modulation of reach variability by success or failure, it did modulate the learning of reward contingencies in the probabilistic learning task. The effects of aging and Parkinson’s disease on reinforcement tasks is thus different even though older-old adults also demonstrate better learning for avoiding choices with low probability of reward compared to older adults ten years younger (Frank & Kong, 2008). Similar results were obtained by Simon and colleagues (2010) using age groups similar to the ones in our study. Yet, we could not replicate the latter effect as older adults in our study were worse at avoiding low-reward conditions compared to young adults.

Given that reward-driven effects are modulated by age (e.g., Düzel et al., 2010; Frank et al., 2004), we expected that the reward-driven increase in implicit adaptation would relate to reward-driven effects observed in typical reward tasks. On the contrary, and in line with Al-Fawakhiri et al. (2023), we failed to find such relationship; both the probabilistic selection task and the controlled probability of reward showed results that appeared unrelated to the implicit adaptation outcome.

Our findings from the different tasks highlight a stark contrast between studies in the motor domain and studies in the cognitive domain where many studies found an age effect even though this age effect might be sensitive to the task and to the context. In the Iowa Gambling task, older adults appear more sensitive to positive reward than younger adults (Bauer et al., 2013). In probabilistic learning tasks or in decision-making tasks, there appear to be clear age-related differences in the sensitivity to positive or negative reinforcement signals (Eppinger et al., 2011). In general, there is a strong tendency for older adults to be particularly sensitive to recent negative outcomes (Worthy et al., 2015). Yet, some studies found that aging does not impair simple reinforcement learning tasks in a three-arm bandit task (Daniel et al., 2020) where attentional demands are limited.

### Aging does not Affect Implicit Motor Adaptation

From our main goal, our results were also clear in pointing that the influence of task outcome and the amount of implicit adaptation are independent of age. The lack of differences between young and older adults in implicit sensory motor adaptation corroborates several recent studies (Cisneros et al., 2024; Hermans et al., 2025; Van De Plas & Orban de Xivry, 2026; Vandevoorde & Orban de Xivry, 2019, 2020, 2021). Two meta-analyses, one using the after-effect as a marker of implicit adaptation and one focusing solely on the task-irrelevant clamped feedback task used in this study, found an enhancement in implicit adaptation with aging (Cohen’s *d* = 0.4, 0.34 and 0.33 for Cisneros et al., 2024, Van De Plas & Orban de Xivry, 2026, and de Witte et al. 2026, respectively). Yet, in both our experiments, the amount of adaptation appears slightly lower in older adults than in young adults. The major difference with the above-mentioned studies is that we used a very small perturbation angle (1.75°) while previous studies used perturbation angles above 30°. There is a need for further research to test the effect of perturbation size to understand fully how aging affects motor adaptation.

## Conclusion

Taken together, our results indicate that the influence of aging on outcome-dependent processes is domain-specific. In the motor domain, reward and outcome signals modulate behavior similarly across age groups, suggesting that feedback-processing mechanisms are largely preserved. In contrast, cognitive tasks involving probabilistic learning show age-related declines, consistent with previous reports of altered reward sensitivity in older adults. Overall, these findings challenge the notion of a uniform age-related decline in reward processing and instead support a more differentiated model in which motor and cognitive systems are differentially affected by aging.

## Acknowledgement

Internal Funds of the KU Leuven (C14/17/115) supported this study. The funder had no role in study design, data collection and analysis, decision to publish, or preparation of the manuscript.

